# Acute sleep deprivation reduces fear memories in male and female mice

**DOI:** 10.1101/2024.01.30.577985

**Authors:** Allison R. Foilb, Elisa M. Taylor-Yeremeeva, Brett D. Schmidt, Kerry J. Ressler, William A. Carlezon

**Author notes:** **Corresponding Author:** William A. Carlezon, Jr., Ph.D., Department of Psychiatry, McLean Hospital, 115 Mill Street, Belmont, MA 02478, USA.

## Abstract

Sleep problems are a prominent feature of mental health conditions including post-traumatic stress disorder (PTSD). Despite its potential importance, the role of sleep in the development of and/or recovery from trauma-related illnesses is not understood. Interestingly, there are reports that sleep deprivation immediately after a traumatic experience can reduce fear memories, an effect that could be utilized therapeutically in humans. While the mechanisms of this effect are not completely understood, one possible explanation for these findings is that immediate sleep deprivation interferes with consolidation of fear memories, rendering them weaker and more sensitive to intervention. Here, we allowed fear-conditioned mice to sleep immediately after fear conditioning during a time frame (18 hr) that includes and extends beyond periods typically associated with memory consolidation before subjecting them to 6 hr of sleep deprivation. Mice deprived of sleep with this delayed regimen showed dramatic reductions in fear during tests conducted immediately after sleep deprivation, as well as 24 hr later. This sleep deprivation regimen also increased levels of mRNA encoding brain-derived neurotrophic factor (BDNF), a molecule implicated in neuroplasticity, in the basolateral amygdala (BLA), a brain area implicated in fear and its extinction. These findings raise the possibility that the effects of our delayed sleep deprivation regimen are not due to disruption of memory consolidation, but instead are caused by BDNF-mediated neuroadaptations within the BLA that actively suppress expression of fear. Treatments that safely reduce expression of fear memories would have considerable therapeutic potential in the treatment of conditions triggered by trauma.

## INTRODUCTION

Fear- and trauma-related learning and memory alter—and are altered by—sleep. Abnormal sleep is a characteristic feature of many fear- and trauma-related disorders. Sleep-related features of Generalized Anxiety Disorder (GAD) and Post-Traumatic Stress Disorder (PTSD) include nightmares, difficulty falling or staying asleep, and restless unsatisfying sleep among their diagnostic criteria^1^. Additionally, these disorders appear to have a reciprocal relationship with sleep, whereby sleep problems worsen symptom severity; indeed, the consequences of insufficient sleep on mental health-related domains including mood and cognitive function are well documented^2–5^. The ability to sleep after experiencing a traumatic event (e.g., an assault, accident, combat mission, mass casualty incident) is often compromised due to various situational parameters, such as the need for medical or police intervention, the need to remain on duty, or subsequent trauma-related insomnia. Interestingly, there are reports that sleep deprivation immediately after a traumatic experience can reduce fear memories, an effect that could be utilized therapeutically in humans^6,7^. Despite its potential importance, however, the role of sleep in the development of and/or recovery from trauma-related illnesses is not fully understood.

In research settings, Pavlovian fear conditioning is commonly used as a model to study threat- and trauma-related human experiences^8,9^. Studies of sleep deprivation on fear memory frequently report memory deficits, but typically focus on the time immediately after trauma, which prevents sleep-based interventions from benefiting clinical populations in most instances^7,10–12^. When sleep is restricted immediately preceding or following fear conditioning, impairments in contextual memory are observed^7,10^. Studies in mice enable exquisite resolution of this effect, showing reductions the expression of fear memories if sleep deprivation occurs during the 0-5 hr period after Fear Conditioning, but no effect if delayed until the 5-10 hr period^7^. Depriving rodents specifically of rapid eye movement (REM) sleep produces similar results, including impaired contextual and cued fear memory as well as impairments in induction of long-term potentiation (LTP)^11,12^. The mechanism by which sleep deprivation immediately following fear conditioning impairs fear memory is thought to involve disruption of memory consolidation. Seminal findings in electrophysiology and animal behavior show that prevention of consolidation through protein synthesis or transcription inhibition in immediate hours after learning or LTP induction prevents formation of the memory, or persistence of LTP^13–16^. Although there is also some evidence that additional (remnant) consolidation occurs in waves requiring a longer time frame (∼12 hr)^17,18^, it is generally accepted that synaptic consolidation—which is required for a memory to last 24 hr and beyond—is complete within a few hours of learning^13^. There is also evidence that sleep disruption can itself serve as type of stress^19,20^, and stress is known to impair or enhance memory function depending on the severity and duration^21–23^. Stress is also known to induce neuroplasticity in brain cells and circuits that regulate the development and expression of fear-related behaviors, including the amygdala^9,24^. Regardless of mechanism, the possibility that early (immediate) sleep deprivation could be used therapeutically to reduce the formation and expression of traumatic memories has important—and potentially exciting—implications for mental health. Importantly, however, this approach would require rapid and highly organized interventions beginning within the minutes immediately following trauma exposure, since the presumed mechanism of action would be preventing the initial formation of fear memories rather than reducing or eliminating already-stabilized fear memories. Approaches that enable intervention at more distal time points—days, weeks, months, or years after trauma—would have a transformational impact on the treatment of conditions like PTSD.

The effects of sleep deprivation at distal time points on fear memories has not been comprehensively characterized. Here, we examined in mice the effects of a delayed, next-day regimen of sleep deprivation of the expression of fear. Specifically, we allowed fear-conditioned mice to sleep immediately after fear conditioning during a time frame (18 hr) that includes and extends beyond periods typically associated with memory consolidation before subjecting them the next day to 6 hr of sleep deprivation. We used a method of sleep deprivation (gentle stimulation) that does not produce stress responses in mice, to avoid adding an additional (secondary) stressor that could confound interpretation of our findings. In parallel, we examined the effects of our sleep deprivation regimen on expression of mRNA encoding brain-derived neurotropic factor (BDNF), molecule implicated in neuroplasticity, in brain areas implicated in the development, expression, and extinction of fear-related behaviors^17,25,26^. Given the well-characterized sex differences in baseline sleep parameters^27–31^, interactions between sleep and gonadal hormones^28,30,32–36^, and the prevalence of trauma-related illnesses such as PTSD^37–44^, we designed the studies to enable qualitative and quantitative comparisons between males and females. Our findings show that delayed sleep deprivation can reduce expression of conditioned fear in both sexes, and raise the possibility that the effects of this regimen are not due to disruption of memory consolidation but instead caused by BDNF-mediated neuroadaptations within the BLA that actively suppress expression of fear.

## METHODS

### Subjects

Adult (6-8 weeks) male and female C57BL/6 mice (Jackson Laboratories, Bar Harbor, ME) were maintained on a 12-hour light/dark cycle (07:00 On-19:00 Off) in a temperature-controlled vivarium. Mice were singly-housed in standard plastic cages with ad libitum food and water, and provided one week of acclimation to the vivarium prior to experiments. Mice that underwent surgery were given 2 weeks for recovery before testing. Experiments included roughly equivalent numbers of male and female mice (except where noted), and all procedures were approved by McLean Hospital Institutional Animal Care and Use Committee and performed in accordance with the National Institutes of Health’s (NIH) Guide for the Care and Use of Animals.

### Sleep Deprivation: Gentle Stimulation and Sweeper Bar

In initial studies, we compared two methods of inducing a 6-hr period of sleep deprivation (gentle stimulation and sweeper bar). Both procedures started at lights-on (07:00), with one mouse per holding cage. For the gentle stimulation method, the mice were monitored for the entire 6-hr period. The procedure began by moving the to a new cage with new nesting materials. Once a nest was made, the mice were prevented from sleeping by introducing novel objects to encourage voluntary activity, as well as gently tapping the holding cage or touching them with a soft brush on the hindquarters. In contrast, the sweeper bar method was automated and involved a sleep fragmentation apparatus (Lafayette Instrument; Lafayette, IN, USA). The apparatus closely resembled the holding cage used for the gentle stimulation method, but was outfitted with a sweeper bar that continuously moved in alternating 7.5-sec cycles across the bottom of the cage. Mice actively avoid contact with the bar, which cycles at a frequency that prevents sleep onset. In light of differences in the stress response caused by these methods (see **RESULTS**), the gentle stimulation method was selected for use in the fear conditioning studies.

### Corticosterone ELISA

To determine if the gentle stimulation and sweeper bar methods produce a stress response, we examined their effects on circulating corticosterone (CORT). Immediately following sleep deprivation, mice exposed to each method were sacrificed by rapid decapitation and trunk blood was collected and centrifuged to obtain plasma. Corticosterone (CORT) levels in plasma were quantified using enzyme-linked immunosorbent assay (ELISA) kits (Enzo Life Science, Ann Arbor, MI, USA) according to manufacturer instructions.

### Fear Conditioning and Extinction Paradigms

All behavioral tests were performed 6 hours after lights on, so that sleep deprivation (on Day 4) would not shift the normal time of testing. As described previously^8,45^, mice were habituated to a fear conditioning chamber (Context A) for 15 minutes on two consecutive days (Days 1-2). Context A consisted of a free-standing chamber (no external sound-attenuating chamber) with a shocker grid floor, house lights on, and lightly scented with quatricide (used as a cleaner). For Fear Conditioning (Day 3), mice were placed in Context A and received 5 conditioned stimulus (CS)-unconditioned stimulus (US) pairings: the CS was a 30-sec, 6000-Hz, 75-dB tone, and the US was a 1-sec, 0.7-mA footshock. Tone-shock trials were presented on a variable ITI, ranging from 1-3 min, and began after an initial 3-min acclimation to the context. The mice were then returned to their home cages and left undisturbed until the next day. The next morning (Day 4), at the start of lights-on cycle—i.e., 18 hr after fear conditioning—half of the mice underwent gentle stimulation sleep deprivation for 6 hours. Control mice were left undisturbed. Immediately after this regimen, mice were tested for Fear Recall in Context B, which was a novel chamber inside an external sound-attenuating chamber, with smooth black plastic flooring, house lights off, and lightly scented with 70% ethanol (used as a cleaner). Fear Recall tests consisted of a 2-min acclimation period, followed by 15 CS (tone) presentations with a 90-sec ITI. For Extinction Recall (Day 5), the Fear Recall procedure was repeated. The primary endpoint for Fear Conditioning tests was freezing behavior, which was quantified using FreezeFrame software (Coulbourn) with freezing thresholds set by a trained observer unaware of treatment conditions.

### Sleep Transmitter Surgery

In experiments designed specifically to examine how fear conditioning affects sleep patterns, mice were implanted with wireless transmitters (HD-X02; Data Sciences International [DSI], St. Paul, MN) to enable continuous collection of EEG (electroencephalography) and EMG (electromyography) data, as described^31,46–48^. Mice were anesthetized via intraperitoneal (IP) injections of 100 mg/kg ketamine/10 mg/kg xylazine mixed in saline. EEG leads were attached to the skill with screws in contact with dura, and secured in place using dental cement. Screws were placed over the frontal lobe (+1mm anterior/posterior, +1mm medial/lateral) and over the contralateral parietal lobe (-3mm anterior/posterior, -3mm medial/lateral). EMG wires were threaded through the trapezius muscle and skin was sutured to close the remaining incision. Antibiotic ointment was applied to the sutured incision, antibiotic (sulfamethoxazole and trimethoprim) was provided in water, and ketofen was given subcutaneously as an analgesic (5.0 mg/kg). After surgery, mice were given 2 weeks to recover prior to the start of baseline sleep recordings that preceded fear conditioning procedures.

### Physiological Recordings

Mice implanted with sleep transmitters were housed in standard plastic cages placed on top of receiver platforms (RPC-1; DSI) for wireless data collection, as previously described^31,46–48^. Collection of EEG and EMG data occurred continuously except when the mice were in the fear conditioning apparatus, enabling analyses before and after habituation sessions and for 24 hours after fear conditioning. Data from habituation Day 2 were used as the baseline for comparison of vigilance states before and after fear conditioning. Vigilance states were quantified using software Neuroscore (DSI) by a trained scorer unaware of treatment conditions.

### Quantitative polymerase chain reaction (qPCR) to quantify mRNA encoding BDNF

In light of previous work implicating BDNF in the development, expression, and extinction of conditioned fear^25,26,49,50^, we examined the effects of our 6-hr gentle stimulation-induced sleep deprivation regimen on BDNF mRNA in brain areas previously implicated in fear and trauma memory processing, including the basolateral amygdala (BLA), medial prefrontal cortex (PFC), bed nucleus of the stria terminalis (BNST), and dorsal hippocampus (HIP)^9,51^. These studies were designed to examine gene expression immediately following sleep deprivation (without the confounding effects of Fear Recall/Extinction testing). Mice were sacrificed by rapid decapitation immediately after 6 hours of sleep deprivation or uninterrupted sleep. Brains were flash frozen using 2-methylbutane and stored in a -80 degree freezer. Brains were sectioned on a cryostat to enable precision dissections (via 0.8mm tissue punches). RNA was extracted with RNeasy Micro Kit (QIAGEN, Germantown, MD, USA) according to manufacturer instructions, cDNA was synthesized with iScript cDNA Synthesis Kit (BIO-RAD, Hercules, CA, USA) and quantitative polymerase chain reaction (qPCR) was performed with iQ SYBR green supermix (BIO-RAD) on MyiQ Single Color Real-Time PCR Detection System (BIO-RAD) with coordinating software. Primers were acquired through Integrated DNA Technologies, Inc. (IDT; Coralville, IA, USA) (see Table 1). BDNF expression was quantified using the Pfaffel method relative to housekeeping gene *7SK*^52^. Values were then normalized to the control group and normalized values were used for statistical analyses. Analyses were performed on the same results as males and females combined, normalized to male and female controls, and as males and females separately, with each sex normalized to their same-sex controls to facilitate within-sex comparisons.

**Table 1.**
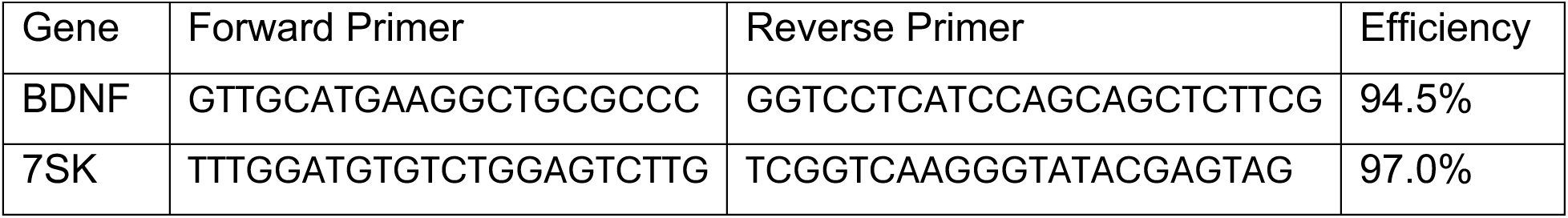
qPCR Primers. Forward and reverse primer sequences, and efficiency values used for Pfaffl calculations.

### Statistical Analyses

Data analyses were performed used GraphPad Prism 9 with significance set to *P*<0.05. Outliers were determined using ROUT outlier detection test (Q=1) and outlier exclusions are noted in Results. Behavioral data were measured as percent freezing for the duration of the cue presentation or baseline. Two-way ANOVA (Sleep Condition x Trial/Trial Block) with repeated measures was used to determine main effects, Tukey’s post-hoc tests were used to compare between trials or trial blocks and Bonferroni’s multiple comparisons was used to compare freezing between sleep conditions. Two-tailed unpaired t-tests were used for analyses of CORT levels and BDNF expression. Two-tailed, paired t-tests were used to analyze changes in vigilance states after fear conditioning relative to baseline sleep.

## RESULTS

### Sleep Deprivation by the Gentle Stimulation and Sweeper Bar Methods Differentially Affect CORT

To determine if our methods for inducing sleep deprivation cause stress responses in mice, we measured differences in plasma CORT in mice sacrificed immediately after each 6-hr regimen. For the gentle stimulation method, when data from the sexes were combined, there were no overall differences in CORT levels between mice that had received sleep deprivation (N=16; 8 males, 8 females) or undisturbed controls (N=18; 8 males, 10 females) (t(32)=1.39, not significant [ns]) (**Fig 1A**). Likewise, there were no differences between conditions when males (t(14)=1.87, ns) and females (t(16)=0.36, ns) were analyzed separately. These data suggest that nether male nor female mice perceive the gentle stimulation method of sleep deprivation as stressful. In contrast, for the sweeper bar method, when data from the sexes were combined, CORT levels were significantly higher in mice that had received sleep deprivation (N=18; 9 males, 9 females) compared to controls (N=17; 9 males, 8 females) (t(33)=3.99, *P*=0.0003) (**Fig 1B**). This effect was also seen when males (t(16)=3.14, *P*=0.0064) and females (t(15)=2.4, *P*=0.03) were analyzed separately. These data suggest that both male and female mice perceive the sweeper bar method of sleep deprivation as stressful. In light of these data, we selected the gentle stimulation method of inducing sleep deprivation for the Fear Conditioning studies, with the justification that use of a stress-free approach in studies of trauma would provide a clearer signal and enable more straightforward data interpretation than would be possible by adding an additional stressor.

**Figure 1.**
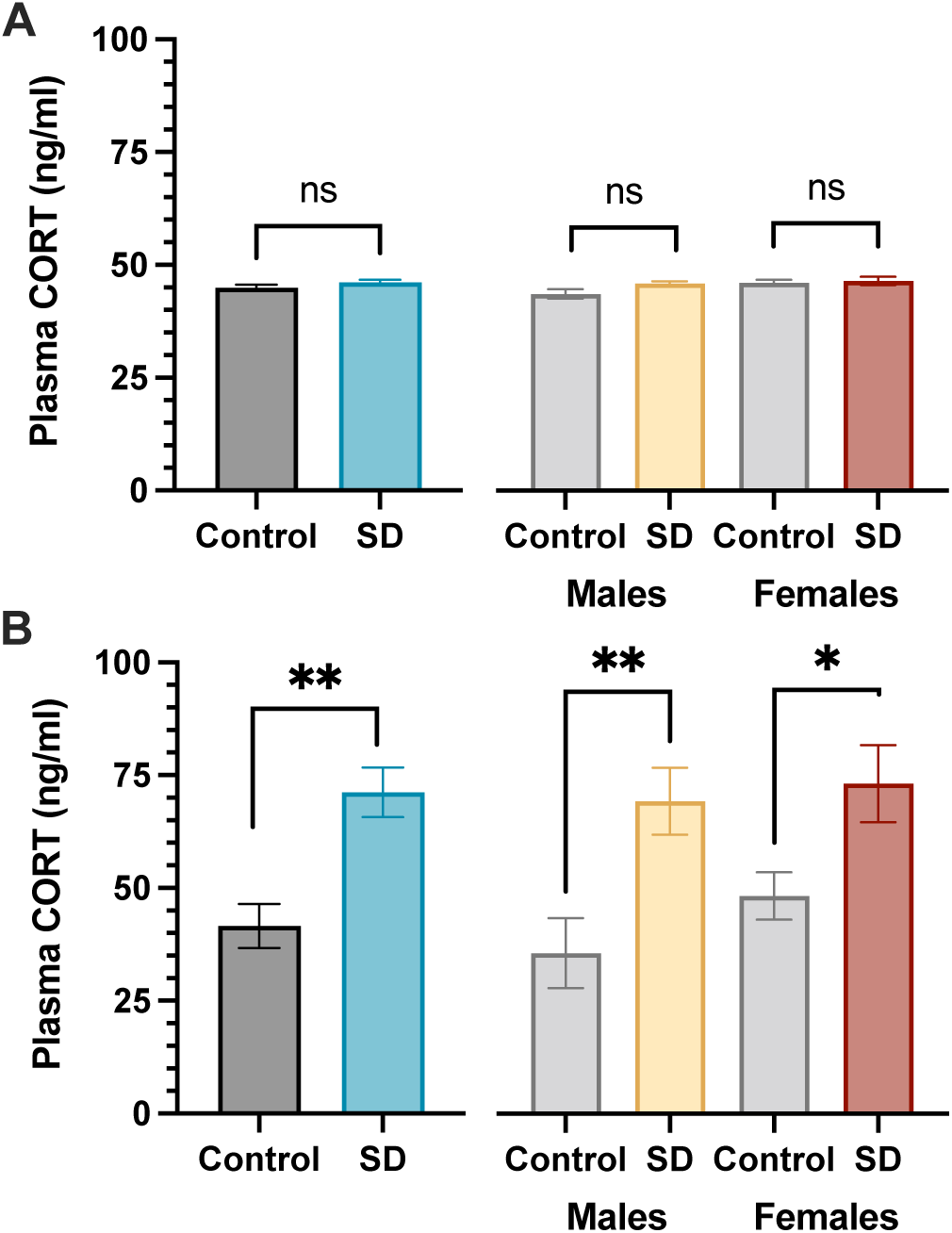
Plasma CORT after sleep deprivation. (A) For the gentle stimulation method, CORT levels in sleep-deprived mice did not differ from those in control (undisturbed) mice when data from the sexes were combined (left) or analyzed separately (right). (B) For the sweeper bar method, CORT levels were significantly higher in sleep-deprived mice compared to controls when data from the sexes were combined (left) or analyzed separately (right). N’s=8-10/sex/group, \**P*<0.05, \*\**P*<0.01, ns=not significant, t-tests.

### Sleep Deprivation Prior to Fear Recall Tests Reduces Expression of Conditioned Fear

As described previously^8,45^, our Fear Conditioning regimen (**Fig. 2A**) produced progressive increases in freezing behavior, our operational measure of fear. During Fear Conditioning (Day 3)—but prior to sleep deprivation—mice did not differ by treatment assignment (n=14/condition; 7M, 7F), and freezing at each trial significantly increased with tone-shock pairings (**Fig 2B**): as would be expected before the sleep deprivation regimen, a 2-way ANOVA revealed a main effect of Trials (F(5,130)=34.69, *P*<0.0001), but no main effect of Sleep Condition (F(1,26)=0.16, ns) and no Trial x Sleep Condition interaction (F(5,130)=0.21, ns). Post hoc comparisons (Tukey’s tests) revealed significant increases in freezing behavior during trials 3-5 when compared to baseline (BL) (*P*’s<0.0001) or trials 1-2 (*P*’s<0.001). In addition, freezing during trial 5 was significantly higher than during trial 3 (*P*=0.02). A similar pattern of effects was seen when the sexes were analyzed separately: there was a significant main effect of Trials in males (F(5,60)=16.14, *P*<0.0001) (**Fig 2C**) and females (F(5,60)=16.89, *P*<0.0001 (**Fig 2D**), and no main effect of Sleep Condition or interaction. In both males and females, post hoc tests revealed significant increases in freezing during trials 3-5 when compared to BL (*P*’s<0.001) and the first trial (*P*’s<0.01). In males, freezing was also increased during trials 3-5 compared to trial 2 (*P*’s<0.01), whereas in females, freezing during trials 4-5 was increased compared to trial 2 (*P*’s<0.01). These data indicate that our Fear Conditioning regimen produces comparable levels of fear in male and female mice.

**Figure 2:**
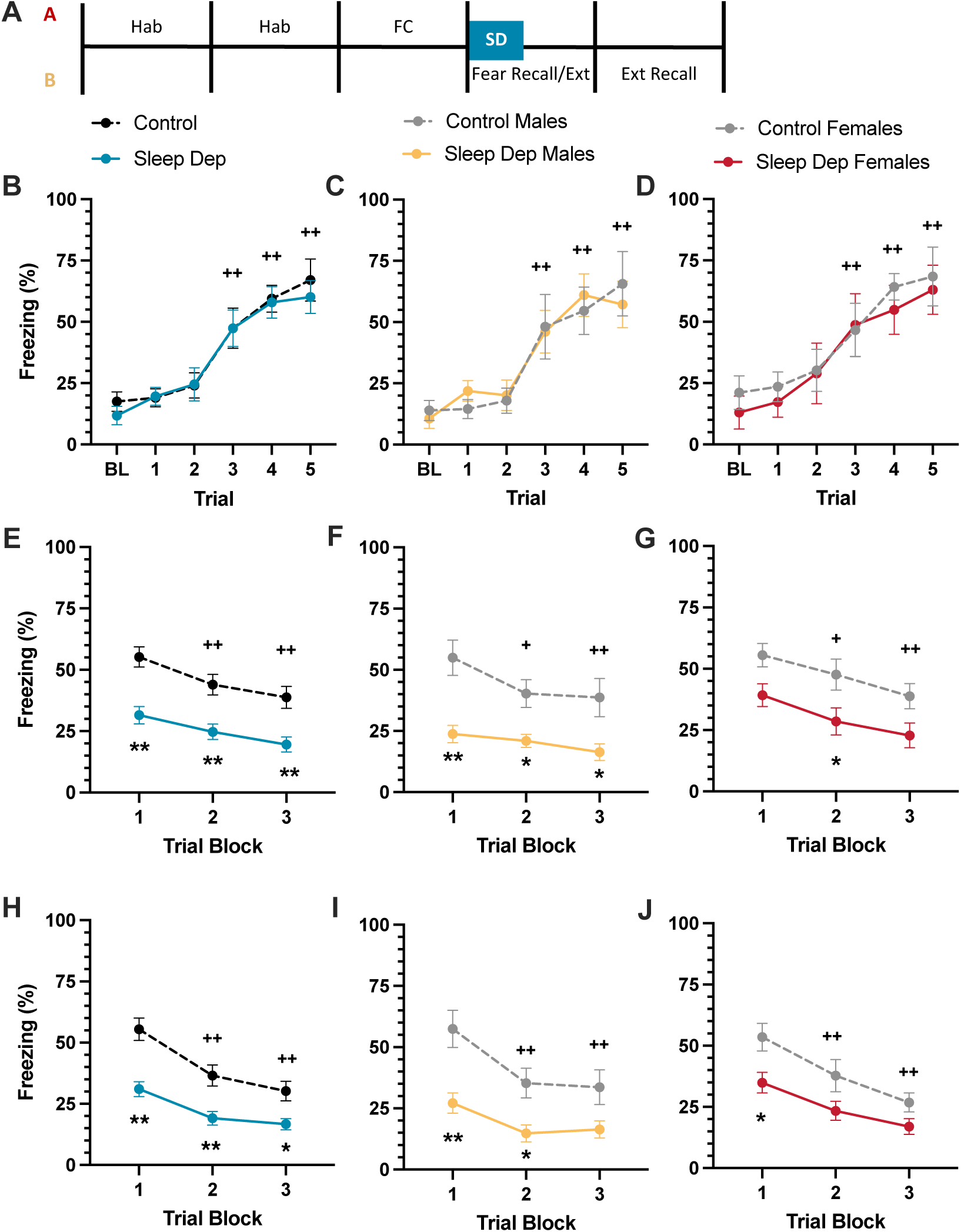
Effects of “delayed” sleep deprivation on fear responses in male and female mice. (A) Schematic of the experimental (“A-B-B”) design. (B) Fear conditioning develops with repeated pairing of the CS (tone) and UCS (footshock), reflected by progressive increases in freezing, in groups where sexes are combined or when data from (C) males or (D) females are analyzed separately. (E) In Fear Recall tests, fear responses diminish progressively over trials, but sleep-deprived mice showed reduced responses immediately, even without extinction trials. This pattern was seen when sexes are combined or data from (F) males or (G) females are analyzed separately. (H) In Extinction Recall tests, sleep-deprived mice continued to show reduced responses, even without further extinction trials, helping to rule out non-specific effects of sleep deprivation in the Fear Recall tests on the previous day. This pattern was seen when sexes are combined or (I) males or (J) females are analyzed separately. Combined N’s=14, individual N’s=7/sex, ++P<0.01, main effect of trials, Tukey’s tests; *P<0.05, **P<0.01 between-group differences, Bonferroni’s tests.

Immediately after our 6-hr sleep deprivation regimen (Day 4), mice underwent Fear Recall testing—which also serves as Extinction training—in response to 15 presentations of the CS (tone) without the US (footshock) in Context B. For clarity, data were consolidated into 3-trial blocks of 5 cue presentations each (**Fig 2E**). A 2-way ANOVA of data with the sexes combined revealed main effects of Trial Block (F(2,52)=20.45, *P*<0.0001) and Sleep Condition (F(1,26)=19.46, *P*=0.0002), but no Trial Block x Sleep Condition interaction (F(2, 52)=0.68, ns). Post hoc comparisons (Tukey’s tests) of Trial Blocks revealed significant reductions in freezing during trial blocks 2-3 compared to trial block 1 (*P*’s<0.001), indicating that fear to the cue had extinguished across the test session. Subsequent Bonferroni’s multiple comparison tests revealed significant reductions in freezing in sleep-deprived mice during trial block 1 (*P*<0.0001) and 2-3 (*P*’s=0.0018). A similar pattern of effects was seen when the sexes were analyzed separately. In males, there were also main effects of Trial Block (F(2,24)=8.84, *P*=0.0013) and Sleep Condition (F(1,12)=12.52, *P*=0.004), with reduced freezing during trial blocks 2 (*P*=0.02) and 3 (*P*=0.0013), and lower levels of freezing in sleep-deprived mice compared to controls during all trial blocks (*P*’s<0.05) (**Fig 2F**). In females, there were also main effects of Trial Block (F(2,24)=11.8, *P*=0.0003) and Sleep Condition (F(1,12)=7.39, *P*=0.019) (**Fig 2G**).

Similar to males, females also displayed significantly reduced freezing in trial blocks 2 (*P*=0.031) and 3 (*P*=0.0002) compared to trial block 1, and lower levels of freezing in sleep-deprived mice compared to controls on trial block 2 (*P*=0.044). Importantly, no main effect of sleep condition was seen during inter-trial intervals (F(1,26)=0.88, ns), suggesting that sleep deprivation did not produce non-specific effects on locomotor activity that would be incompatible with freezing behavior (not shown). These data indicate that sleep deprivation produces immediate reductions in freezing, even in the absence of extinction training, with minimal qualitative differences between sexes.

On the following day (Day 5), mice were returned to Context B for Extinction Recall tests, which were performed an analyzed exactly as the Fear Recall tests on the previous day. Consistent with the Fear Recall tests, a 2-way ANOVA revealed main effects of Trial Block (F(2,52)=42.32, *P*<0.0001) and Sleep Condition (F(1,26)=17.94, *P*=0.0003), but no Trial Block x Sleep Condition interaction (F(2,52)=2.99, ns) (**Fig 2H**). Again, post hoc comparisons (Tukey’s tests) of Trial Blocks revealed significant reductions in freezing during trial blocks 2-3 compared to trial block 1 (*P*’s<0.0001), indicating further extinction of fear responses to the cue. Subsequent Bonferroni’s multiple comparisons again revealed significant reductions in freezing in sleep-deprived mice during all trial blocks: 1 (*P*<0.0001), 2 (*P*=0.003), and 3 (*P*=0.03). A similar pattern of effects was seen when the sexes were analyzed separately. In males, there were also main effects of Trial Block (F(2,24)=18.41, *P*<0.0001) and Sleep Condition (F(1,12)=10.83, *P*=0.0064), with reduced freezing during trial blocks 2-3 (*P*’s<0.0001), and lower levels of freezing in sleep-deprived mice compared to controls during Trial Block 1 (*P*=0.0014) and 2 (*P*=0.039) (**Fig 2I**). In females, there were also main effects of Trial Block (F(2,24)=26.29, *P*<0.0001) and Sleep Condition (F(1,12)=6.43, *P*=0.026). Similar to males, females also displayed significantly reduced freezing in trial blocks during trial blocks 2-3 (*P*’s<0.0001), and lower levels of freezing in sleep-deprived mice compared to controls on trial block 1 (*P*=0.025). These data indicate that the effects of sleep deprivation on expression of conditioned fear are sustained, while also mitigating concerns that reductions in fear responses during the Fear Recall tests on the previous day reflect a transient artifact of acute sleep deprivation.

### Fear and Sleep Appear Normal at the *18*-hr Post-Fear Conditioning Time Point

In light of the dramatic effect of our sleep deprivation regimen on the expression of conditioned fear, we performed experiments in small numbers of mice to confirm that (a) fear responses are intact at 18 hr and do not differ from those seen at 24 hr, and (b) that sleep patterns are not fundamentally different during the 18 hr after Fear Conditioning. To examine fear responses, we performed our normal Fear Conditioning training in male mice (N=6) but instead performed Fear Recall at 18 hr, and compared these data to those from the mice tested at 24 hr (i.e., **Fig. 2E**). For simplicity, the means of the first 5 trials were compared (**Fig 3A**). A 2-way ANOVA comparing Fear Recall at 18 hr and 24 hr reveal no main effects of Trial (F(4,72)=1.25, ns) or Timepoint (F(1,18)=1.04, ns), nor Trial x Timepoint interaction (F(4,72)=0.3, ns). These results indicate equivalent levels of freezing behavior at 18 hr and 24 hr after Fear Conditioning, ruling out concerns that fear memories are not yet formed at the time we performed our sleep deprivation regimen.

**Figure 3:**
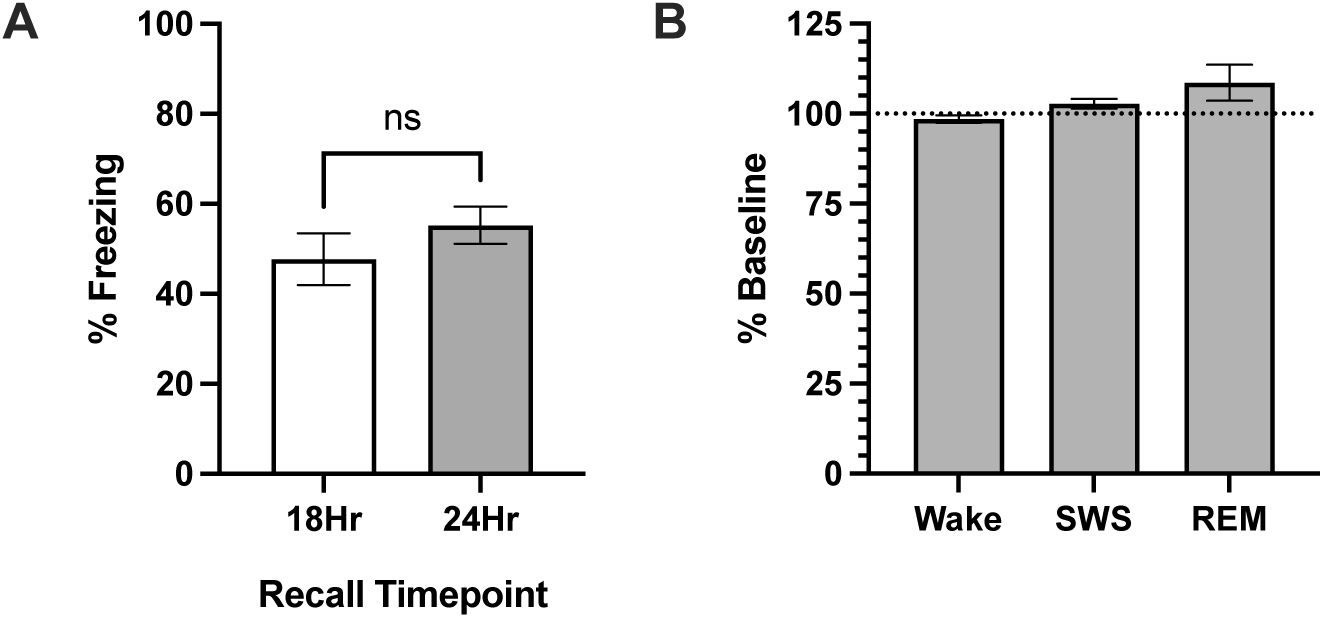
Fear recall and sleep are normal 18 hr after fear conditioning. (A) Fear Recall is similar in tests conducted at 18 hr or 24 hr after fear conditioning. (B) There are no significant changes in sleep architecture during the 18 hours after fear conditioning.

Likewise, we examined sleep architecture during the 18 hr after fear conditioning in a separate cohort of mice (N=29, 15 males, 14 females). Vigilance state durations during the 18 hr after fear conditioning were compared to the same 18 hr time period following context habituation the previous day (i.e., Day 2) (**Fig 3B**). Within-subject analyses (paired t-tests) revealed no changes in duration of wakefulness (Wake; t(28)=1.54, ns), slow wave sleep (SWS; t(28)=1.85, ns), or rapid eye movement sleep (REM; t(28)=0.97, ns). The patterns were similar when males and females were analyzed separately (not shown). These findings indicate that mice sleep normally—without evidence of insomnia—in the hours following Fear Conditioning, a time during which sleep can enhance memory consoliation^43,53^.

### Sleep Deprivation Produces Changes in BDNF Expression

Our sleep deprivation regimen may trigger neuroplastic changes in fear circuits that contribute to suppression of fear responses. Since work from our group and others have implicated BDNF in regulation of fear expression^25,49^, we examined the effects of our 6-hr gentle stimulation-induced sleep deprivation regimen on BDNF mRNA in brain areas previously implicated in fear and trauma memory processing, including the BLA, PFC, HIP, and BNST. These studies were designed to examine gene expression immediately following sleep deprivation, without the confounding effects of Fear Recall/Extinction testing, in males (N=8/condition) and females (N=8/condition). For the BLA, one male (control) and one female (sleep-restricted) were identified as statistical outliers and excluded from analyses. Between-group analysis (t-tests) combining both sexes revealed significant increases in BDNF mRNA (t(28)=3.23, *P*=0.0032) (**Fig 4A**). A similar pattern of results we seen when the sexes were analyzed individually: BDNF mRNA levels were higher following sleep deprivation in males (t(13)=2.36, *P*=0.035) and females (t(13)=2.19, *P*=0.047). For the PFC, one male (control) was excluded as a statistical outlier. Between-group analysis combining both sexes revealed significant increases in BDNF mRNA (t(29)=2.20, *P*=0.036) (**Fig 4B**). However, analysis of the sexes individually revealed a sex-dependent effect: sleep deprivation did not alter BDNF mRNA levels in males (t(13)=0.55, ns), whereas it elevated levels in females (t(14)=2.38, *P*=0.032). For the BNST, between-group analysis combining both sexes revealed no effects of sleep deprivation (t(30)=1.63, ns) (**Fig 4C**). However, analysis of the sexes individually revealed a sex-dependent effect opposite to that seen in the PFC: sleep deprivation elevated BDNF mRNA levels in males (t(14)=3.03, *P*=0.0089), whereas it had no females (t(14)=0.05, ns). For the HIP, one male control was identified as a statistical outlier and excluded. Analyses found no effect of sleep deprivation on BDNF mRNA, regardless of whether sexes were combined (t(29)=0.15, ns) (**Fig 4D**), or if separate analyses were performed for males (t(13)=0.49, ns) and females (t(14) = 0.31, ns). These data demonstrate that our sleep deprivation regimen can trigger increases in BDNF gene expression in brain areas implicated in fear expression, but of the areas examined in this report, only the changes in the BLA were consistent across sexes.

**Figure 4:**
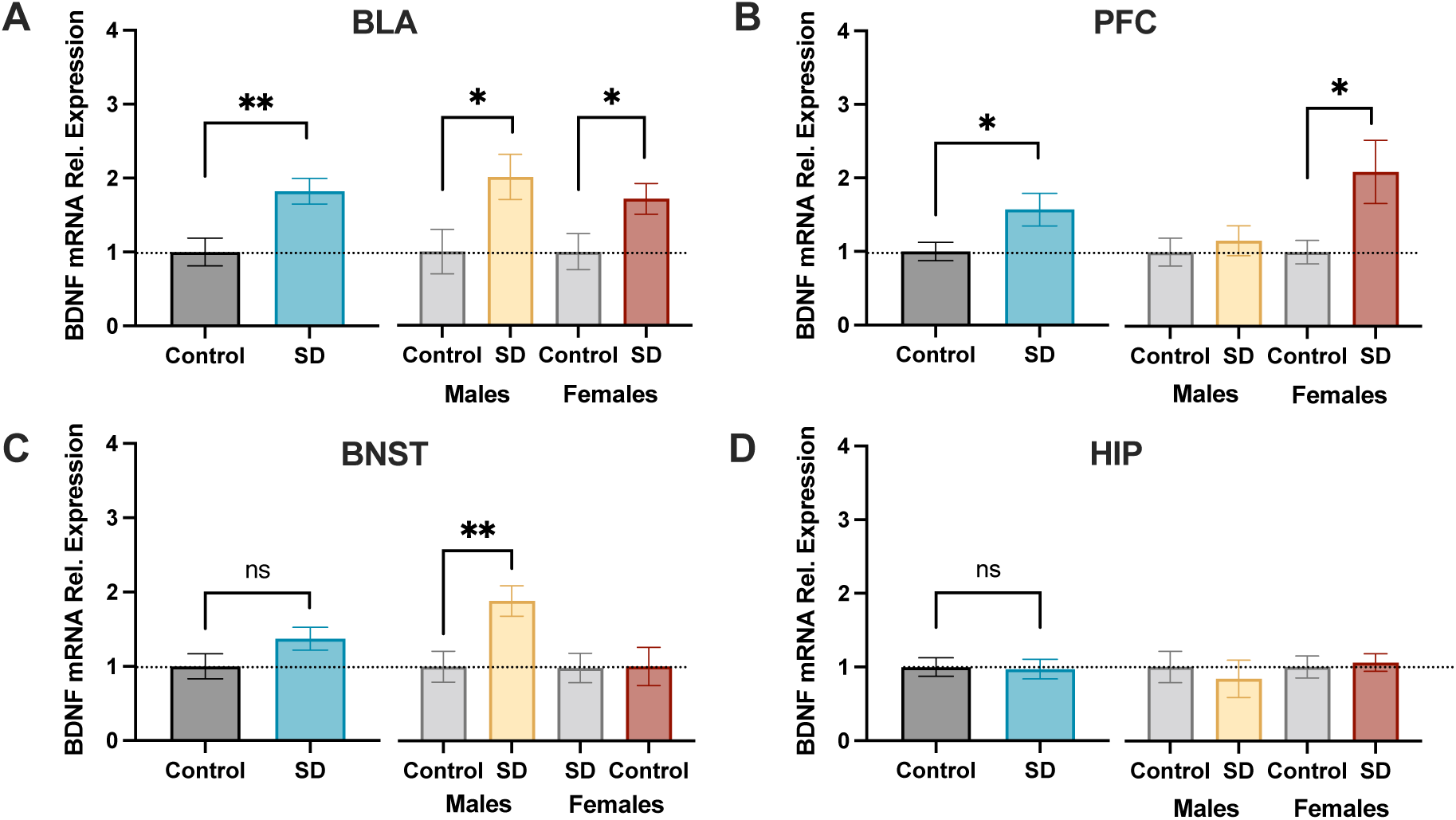
Effects of gentle handling-induced sleep deprivation on expression of mRNA encoding BDNF across numerous brain regions in male and female mice. (A) In the BLA, sleep deprivation produced significant increases in BDNF gene expression when sexes are combined (left) or analyzed separately (right). (B) There were also significant increases in the PFC when sexes were combined, but this effect was carried by females, with no effects in males. (C) In the BNST, increases were seen in males only, with no effects in females alone or when the sexes were combined. (D) There were no effects in the HIP. Combined N’s=16, individual N’s=8/sex, ns=not significant, *P<0.05, **P<0.01, t-tests.

## DISCUSSION

Here we show that a delayed regimen of sleep deprivation can reduce the expression of conditioned fear in mice. A novel and distinguishing feature of these studies is that the sleep deprivation was performed on the day after fear conditioning, following an 18-hr period when the mice were left undisturbed and thus able to sleep normally. While there have been other reports that sleep deprivation can disrupt fear behaviors, in general those have demonstrated that it is necessary to perform the sleep deprivation immediately, beginning within the minutes after fear conditioning^6,7^. The effects were qualitatively and quantitatively similar in similar in males and females, suggesting that they involve mechanisms that are similar across sexes. Importantly, the effects were seen in recall tests conducted immediately following sleep deprivation as well as the day after, ruling out the possibility that is due to a transient, non-specific artifact of the sleep deprivation regimen itself. In addition, we demonstrate that the mice do not perceive the sleep deprivation regimen as stressful, which rules out the possibility that the effects are due to interactions among various stressors. This type of approach may have translational relevance and benefits: the ability to reduce fear at time points that do not require immediate intervention would have considerable therapeutic potential, and if it someday becomes possible to utilize sleep deprivation in traumatized humans for treating conditions like PTSD, procedures that minimize stress would be preferable.

Previous reports describing the effects of sleep deprivation on expression of conditioned fear generally attribute reductions in fear behavior to disruption in memory consolidation. Synaptic consolidation—which is required for a memory to last 24 hr and beyond—is thought to begin immediately following learning and to be complete within a few hours^13^. Studies in mice enable exquisite resolution of this effect, showing reductions the expression of fear memories if sleep deprivation occurs during the 0-5 hr period after Fear Conditioning, but no effect if delayed until the 5-10 hr period^7^. Several observations suggest that the effect of our sleep deprivation effect is not due to disruption of consolidation. Our sleep deprivation regimen began the next day, 18 hr after Fear Conditioning, at a time that includes and extends beyond periods typically associated with memory consolidation. During the 18-hr period, the mice are left undisturbed and allowed to sleep, which is known to enhance memory consolidation^7,43,53^. Using mice implanted with wireless telemetry devices that enable continuous monitoring of sleep architecture, we show that sleep after Fear Conditioning is essentially normal; if anything, there are nominal (though not statistically significant) increases in REM sleep, which in humans has been associated with enhanced consolidation of emotional memories^54^. We also showed in control mice that Fear Recall at the 18-hr time point is equivalent to that seen at the 24-hr time point, suggesting effective consolidation of fear memories. In addition, several observations suggest that our effect is not an artifact of acute sleep deprivation occurring immediately before Fear Recall tests. As one example, we did not observe differences in freezing behavior during inter-trial intervals in Fear Recall tests, ruling out the possibility that sleep deprivation produces non-specific locomotor-activating effects that are incompatible with freezing behavior. Consistency across the Fear Recall and Extinction Recall tests, which are performed 24 hr apart, also suggests that our sleep deprivation regimen produced a sustained effect on fear expression rather than a transient artifact. Importantly, comparison of freezing behavior in control mice (which were left undisturbed during the sleep deprivation regimen) in the Fear Recall and Extinction Recall tests shows intra-session extinction, but limited retention of extinction between sessions. These findings suggest that our protocol produces relatively weak extinction. This is likely due, at least in part, to the use of relatively few Fear Recall test trials, which also serve as extinction trials because they involve repeated presentation of the tone (CS) in the absence of the footshock (US)^55^. The protocol was intentionally designed in this way because in early (pilot) studies that involved more trials and thus were considerably longer in length, we found that some sleep-deprived mice feel asleep during the Fear Recall sessions. Importantly, sleep during sessions can cause apparent increases in freezing—opposite to what we observed in our current report—that are due to sleep-related decreases in spontaneous activity rather than expression of conditioned fear. We did not observe any cases of the mice falling asleep during the Fear Recall or Extinction Recall protocols used here, which showed decreases in freezing in sleep-deprived mice. Future studies may involve different permutations of the current protocols; sleep deprivation regimens that are performed at far longer periods (weeks, months) are a particularly high priority, but these require identification of fear conditioning parameters that engender sustained expression of fear in control groups.

There are also potential explanations for these findings that do not involve disruption of fear memory consolidation. As an example, there are reports that administration of neurotrophic factors can reduce the expression of fear, effectively serving as a “substitute” for extinction training. In a seminal study in rats, it was shown that infusion of BDNF directly into the infralimbic (IL) PFC on the day after Fear Conditioning reduces expression of fear, even without extinction training^49^. Interestingly—and similar to our current findings—complete reductions in fear were seen immediately (in the first Fear Recall trials), as opposed to enhancement of a more prototypical extinction response in which fear responses are seen initially but decay with subsequent CS-only trials. While the neural mechanisms of this effect are not completely understood, there is evidence that BDNF actions in pathways linking the HIP with the IL-PFC are involved^49^. Further data suggest that IL projections to BLA are involved in fear inhibition and extinction^9^. Collectively, the data in those previous reports align with our current finding that our sleep deprivation regimen produces immediate elevations in BNDF mRNA within the BLA. While effects on BDNF were also seen in other brain areas (PFC, BNST), the only the effects in the BLA were consistent in males and females, matching the consistency across sexes in the behavioral (Fear Conditioning) studies. Our finding of increased BDNF gene expression in BLA aligns with previous data supporting a role for BDNF and its receptor (TrkB) within this region in fear behavior and its extinction^9,25,26,50,56^. One intriguing possibility is that BDNF systems are involved in an active process that enables rapid transitions between defensive and exploratory behavior. In this formulation, transitions between high and low fear are triggered by switches in the balance of activity in two distinct populations of BLA neurons, conceptualized as “Fear” and “Extinction” neurons^57^.

Indeed, putative populations of “Fear-On” and “Fear-Off” cells within the BLA have been described^58,59^. Although these neuronal populations are intermingled within the BLA, they receive differential inputs: importantly, Fear-Off neurons are innervated primarily by the mPFC^57^, providing a potential explanation that links the prior work with intra-PFC infusions of BDNF with our current findings^49^. Together, our behavioral and molecular findings provide the basis for studies that explore the possibility that our sleep deprivation regimen recruits BDNF-dependent processes in the BLA that actively suppress fear memories.

In summary, we have discovered a sleep deprivation regimen in mice that can be used the day after Fear Conditioning to reduce expression of fear memories. These findings provide the basis for future work designed to understand the basic molecular, cell-type specific, and circuit mechanisms underlying threat regulation in mammalian amygdala. Furthermore, this work may also provide a basis for new (non-pharmaceutical) therapeutic approaches that could be rapidly translated to the clinic and transformational with respect to the prognosis for people with conditions such as PTSD, even as neural mechanisms are dissected and thoroughly characterized.

